# Circadian rhythms in multiple behaviors depend on sex, neuropeptide signaling, and ambient light

**DOI:** 10.1101/2022.08.18.504454

**Authors:** Lari Rays Wahba, Blanca Perez, KL Nikhil, Erik D. Herzog, Jeff R. Jones

**Affiliations:** Department of Biology, Washington University in St. Louis, St. Louis, MO, USA; Department of Biology, Texas A&M University, College Station, TX, USA

## Abstract

Organisms have evolved circadian (near-24 h) rhythms in behavior to anticipate daily opportunities and challenges such as mating and predation. However, the ethological investigation of circadian behavioral rhythms has been traditionally limited to studying easy-to-measure behaviors at higher temporal resolution or difficult-to-measure behaviors with limited temporal resolution. Our ability to simultaneously record circadian rhythms in multiple behaviors has also been limited by currently available technology. We thus sought to examine eight overt, ethologically-relevant behaviors never before studied simultaneously as a function of time of day: eating, drinking, grooming, rearing, nesting, digging, exploring, and resting. To address the hypothesis that the daily patterning of these behaviors depends on neuropeptide signaling, sex, and ambient light, we used high-throughput machine learning to automatically score millions of video frames of freely-behaving male and female wild-type and vasoactive intestinal peptide (*Vip*)-deficient mice. Automated frame-by-frame predictions of the eight behaviors correlated highly with consensus labels by trained human classifiers. We discovered reliable daily rhythms in many previously unreported behaviors that peaked at stereotyped times of day and persisted in constant darkness. Surprisingly, nesting and digging rhythms differed dramatically in both phase and amplitude between male and female mice. In *Vip*-deficient mice, daily rhythms in most behaviors were low amplitude and peaked earlier in the day in a light:dark cycle, while rhythms in all behaviors peaked randomly throughout the day in constant darkness. We also found that for most behaviors, time budgets predominantly differed by light cycle, but transition probabilities predominantly differed with VIP signaling and by sex. We conclude that machine learning can be used to reveal novel sex, neuropeptide, and light-dependent behaviors at time scales from seconds to days.

## Introduction

Understanding the genetic, neural, and ethological mechanisms that facilitate the temporal organization of behavior is a fundamental goal of fields including neuroscience, ecology, motion science, and circadian biology. One such behavior, locomotor activity, has been widely studied at high temporal resolution using running wheels or infrared beam breaks in animals and wrist actigraphy in humans (Ancoli-Israel et al., 2003; Jud et al., 2005). Other behaviors such as eating and drinking have been measured with optical or electrical sensors (Schwartz and Zimmerman, 1990; Pendergast et al., 2013). However, these behaviors have mostly been studied independently due to the lack of reliable technology for simultaneous behavioral data acquisition. Moreover, determining the temporal structure of multiple complex behaviors typically requires manual scoring by trained observers of recorded videos and is thus infeasible over long timescales. Behaviors that capture the entirety of an individual animal’s daily behavioral repertoire therefore need to be simultaneously measured over multiple days and conditions to better understand how animal behavior changes over time (Garner, 2017).

In mammals, many behaviors are coordinated by the central circadian pacemaker, the suprachiasmatic nucleus (SCN) of the hypothalamus, to anticipate daily environmental challenges such as light and dark, food availability, mating opportunities, and predator avoidance (Yerushalmi and Green, 2009; Hastings et al., 2018). The SCN depends on a population of neurons that produce the neuropeptide vasoactive intestinal peptide (VIP) to maintain synchrony among its cells and with the environment (Aton et al., 2005; Jones et al., 2018; Mazuski et al., 2018; Paul et al., 2020; Todd et al., 2020). Mice genetically deficient for *Vip* or its receptor exhibit disrupted circadian rhythms in numerous physiological processes including glucocorticoid production, metabolism, cardiovascular function, and body temperature (Bechtold et al., 2008; Loh et al., 2008; Schroeder et al., 2011). VIP is also required for normal circadian rhythms in locomotor behavior, as the onset of wheel-running activity in *Vip*-deficient mice housed in a 12 h:12 h light:dark cycle (LD) is about 8 h earlier than in wild-type mice while these rhythms are lost in constant darkness (DD) (Colwell et al., 2003; Aton et al., 2005). However, whether VIP is essential for daily rhythms in other behaviors is unknown.

Sexual dimorphisms are observed in the temporal patterning of numerous physiological processes (Krizo and Mintz, 2014; Joye and Evans, 2022). For example, sex differences have been identified in daily rhythms in glucocorticoid production, cardiovascular function, body temperature, and immune function (Walton et al., 2022). Sex differences have also been observed in wheel-running activity rhythms, although these behavioral differences are much more subtle (Lee et al., 2004; Kuljis et al., 2016). For instance, male mice show a greater precision of wheel-running activity onsets in LD and female mice show a longer wheel-running activity duration in DD (Kuljis et al., 2013). These behavioral differences are likely due to differences in levels of circulating sex hormones and sex hormone receptor expression. However, given the global regulation of multiple behaviors by the circadian system, more work is needed to reveal and understand sex differences in other behavioral rhythms.

Here, we tested the hypothesis that VIP signaling, sex, and light cycle each affect daily patterns of multiple ethologically-relevant behaviors: eating, drinking, grooming, rearing, nesting, digging, exploring, and resting (Garner, 2017). We used DeepEthogram, a general-purpose machine learning classifier, to measure behaviors never before studied simultaneously as a function of time of day in male and female wild-type and *Vip*-deficient mice housed in LD and in DD (Bohnslav et al., 2021). We identified circadian rhythms in several established and novel behaviors with amplitudes and peak times that were strongly dependent on VIP signaling, sex, and light cycle. We conclude that the temporal structure of multiple behaviors depends on neuropeptide signaling, sex, and ambient light, and that machine learning can be used to discover circadian phenotypes that were previously difficult or impossible to observe.

## Results

### Automated classification of behaviors in individual mice at one-second resolution over several days

To validate our model’s performance, we first used it to infer behaviors on a manually-labeled 24 h video of a freely-behaving mouse recorded at one frame per second (86,400 frames; **Supplementary Fig. 1a**). We compared the inferred labels to our manual labels and calculated an F1 score that accounted for both true and false positives and negatives. Our model had an F1 score of ≥ 75% for 7 out of 8 behaviors, and an F1 score of 52% for the “rearing” behavior. To determine how these F1 values compared to chance, we simulated ten randomly-generated ethograms (arrays of behaviors) and compared their F1 scores versus our ground truth manually-labeled frames. Chance F1 values for our simulated random arrays were much smaller than the calculated F1 scores from our model: 14.1 ± 1.8 % (eating, mean ± SEM), 0.4 ± 0.9 % (drinking), 20.3 ± 3.9 % (grooming), 0.4 ± 1.3% (rearing), 4.6 ± 0.7 ± (nesting), 9.9 ± 1.8 % (digging), 7.5 ± 1.3 % (exploring), and 43.8 ± 12.8 % (resting). We also inferred the same video twice through our model and found that reproducibility was high (>90%) between inferences.

**Figure 1.**
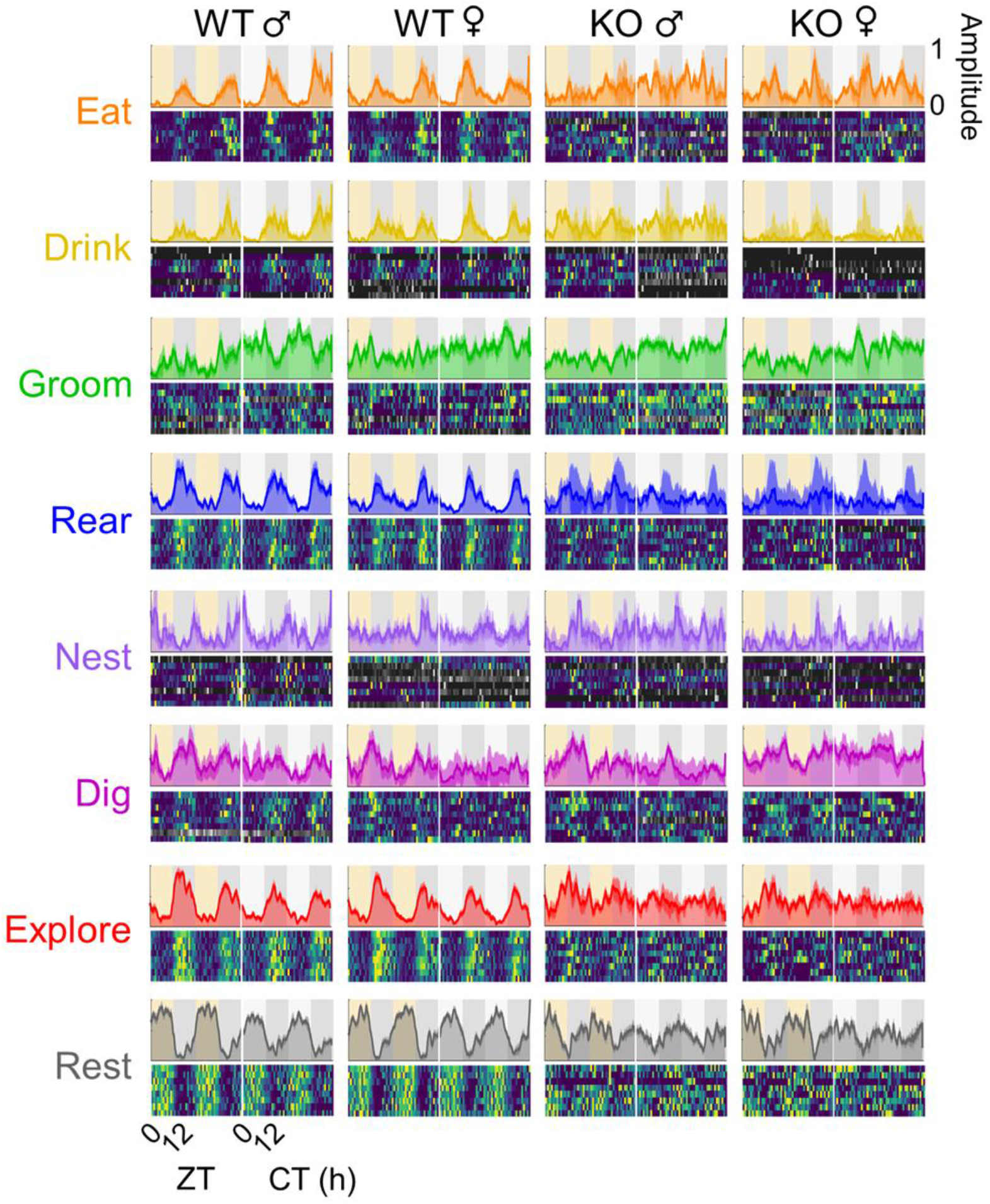
Diurnal and circadian rhythms in multiple behaviors in male and female wild-type and *Vip*-deficient mice. *Top*, ethograms of average behaviors (colored lines and dark colored shading, mean ± SEM) from mice (n = 8 wild-type (WT) male, 8 wild-type female, 8 *Vip*^-/-^ (KO) male, 8 *Vip*^-/-^ female) recorded over 96 h in a 12 h:12 h lightdark cycle (LD, yellow and dark gray background) and in constant darkness (DD, light and dark gray background). Y axes show activity counts smoothed with a 1 h running average and scaled to the maximum for each behavior. *Bottom*, raster plots of hourly activity counts from each mouse in LD and DD with warmer colors indicating greater activity. Activity profiles of individual mice were scored as either circadian (color) or arrhythmic (grayscale). ZT, zeitgeber time. CT, circadian time.

Next, we wanted to determine how well our model performed compared to naive human classifiers. First, three trained human classifiers independently labeled behaviors in four 5 min videos (1,200 frames) of a wild-type mouse. These labels were compared to generate a “reference standard” label based on majority agreement among the trained human classifiers. We then had three naive human classifiers to label the same set of videos and also inferred the same videos using our model. We thus generated one “human label” and “model label” each for the four videos. We compared these labels to the “reference standard” labels and found that the accuracy (defined as the percentage of labeled frames that were identical to the reference standard) of our model did not differ significantly from human classifiers for any of the 8 behaviors analyzed (**Supplementary Fig. 1b**; Two-Way ANOVA, p > 0.500 for each behavior).

Finally, we wanted to compare the performance of our model with another automated behavior analysis program. We used MouseActivity (Zhang et al., 2020) to analyze the position of a mouse and automatically calculate several variables including distance traveled and velocity over time in four video clips previously labeled by our model. To directly compare MouseActivity-predicted behaviors with those of our model, we thresholded the MouseActivity output by defining “exploring” behavior as a minimum path length of 9 mm and “resting” behavior as a frame with no observed movement. We found remarkably high (>80%) agreement between our model’s and MouseActivity’s predictions of whether a given frame contained exploring or resting behavior (**Supplementary Fig. 1c**). Together, these methods of validation demonstrate that our model can reliably and accurately predict behaviors.

### Mice exhibit reliable diurnal and circadian rhythms in behaviors that peak at stereotyped times

Next, we used our validated model to determine how behaviors differ over time with or without VIP signaling, between males and females, and in a light cycle or in constant darkness. We recorded videos at one frame per second of freely-behaving male and female wild-type and Vip-/- mice (n = 32; 8 wild-type and 8 Vip-/- males and females each) for 48 h in LD followed by 48 h in DD. We then used our trained model to predict behaviors. We transformed the resulting binary prediction matrix for each mouse into ethograms depicting the occurrence of each behavior over time for both individual animals and for mice averaged within each genotype and sex (**Fig. 1**) (Lehner, 1998).

We first wanted to determine the temporal sequence of behaviors on the hours-to-days timescale, that is, whether mice exhibited diurnal (in LD) and/or circadian (in DD) rhythms in each recorded behavior. To do this, we first used several independent circadian analysis methods to identify which behaviors were significantly rhythmic in individual mice (**Supplementary Fig. 2**) (Refinetti et al., 2007; Zielinski et al., 2014; Wu et al., 2016). We then used a harmonic regression model to determine the amplitude (the “strength” of the rhythm) and phase (the peak time) of each rhythmic behavior in individual mice. We defined the amplitude of arrhythmic behaviors for each mouse as zero and excluded these behaviors from phase analysis. Finally, we used circular statistics to determine whether mice of a given genotype or sex each exhibited the same entrained and endogenous phase for each rhythmic behavior in LD and in DD, respectively (Berens, 2009).

**Figure 2.**
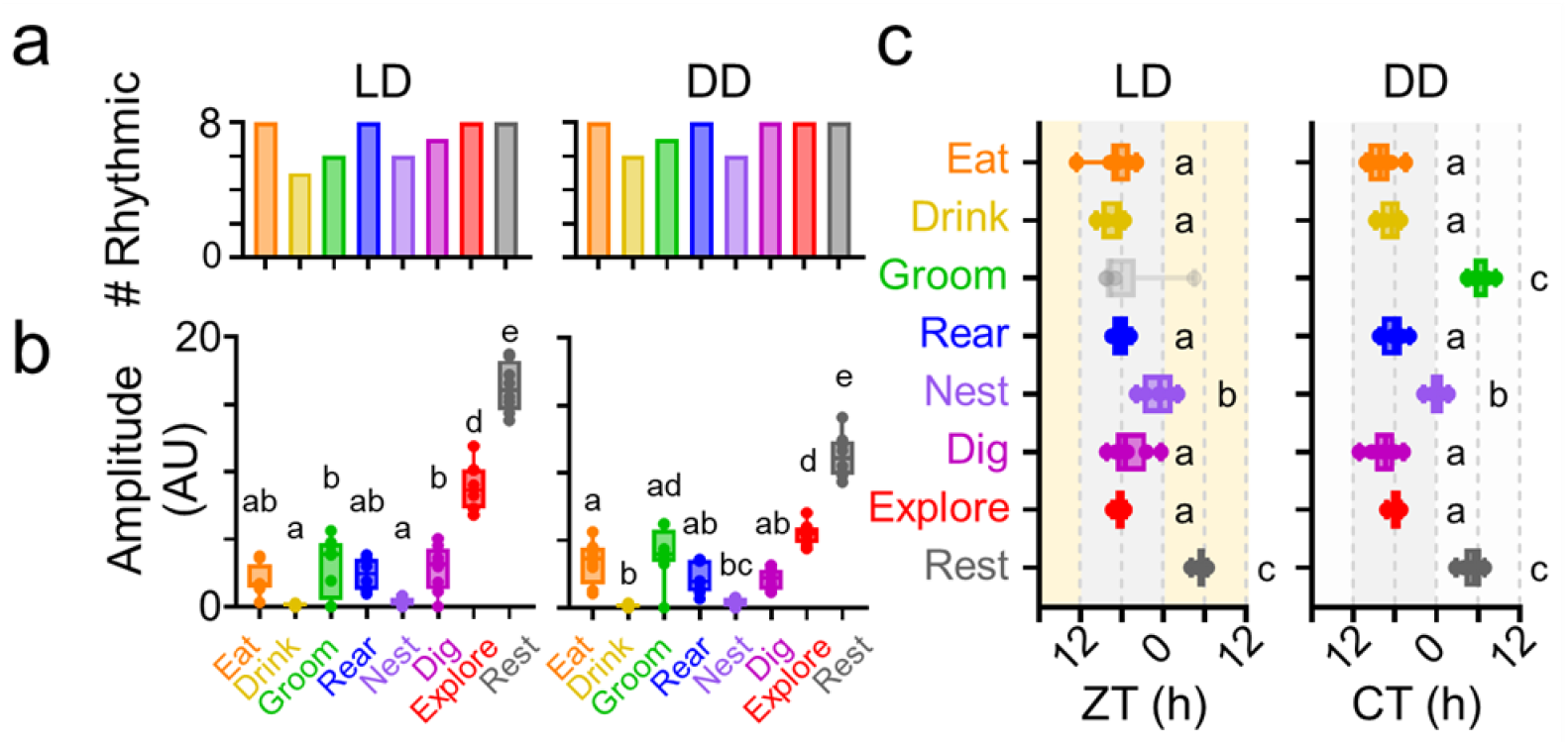
Wild-type male mice exhibit prominent diurnal and circadian rhythms in most behaviors. **a)** Number of individual wild-type male mice (n = 8) recorded in a 12 h: 12 h lightdark cycle (LD) and in constant darkness (DD) scored as rhythmic for each behavior (ARSER, p < 0.05 or less), **b)** Rhythm amplitudes (in arbitrary units) for each behavior in LD and in DD. Boxes indicate 25^th^ to 75^th^ percentiles. Letters on each plot indicate similar amplitudes within each light cycle. (Two-Way Repeated Measures ANOVA, p < 0.05 or less), **c)** Peak times (in hours) for each behavior in individual mice in LD (yellow and dark gray background) and in DD (light and dark gray background). ZT, zeitgeber time. CT, circadian time. Boxes indicate 25^th^ to 75^th^ percentiles. Gray dots and boxes depict rhythms that did not have significant clustering of peak times across mice (Rayleigh test, p > 0.05). Letters on each plot indicate behaviors that peaked at similar times (Watson-Williams tests, p < 0.05 or less). Peak times depicted as linear for visualization.

We found that, consistent with previous reports, most wild-type male mice exhibited prominent daily rhythms (ARSER, p < 0.05) in eating, drinking, exploring, and resting behaviors that persisted in constant darkness (**Fig. 2a**). The amplitudes of exploring and resting behaviors were greater in LD (Three-Way Repeated Measures ANOVA, p < 0.001), but the amplitudes of “maintenance” behavior rhythms (eating and drinking) were greater in DD (p < 0.05 or less; **Figs. 2b, 3**; **Supplementary Fig. 3**) (Garner, 2017). The phase of each behavioral rhythm was highly similar across individual mice (Rayleigh test, p < 0.05) and did not differ between LD and DD (Multi-Way Circular ANOVA, p > 0.05; **Fig. 2c**; **Supplementary Fig. 4**).

**Figure 3.**
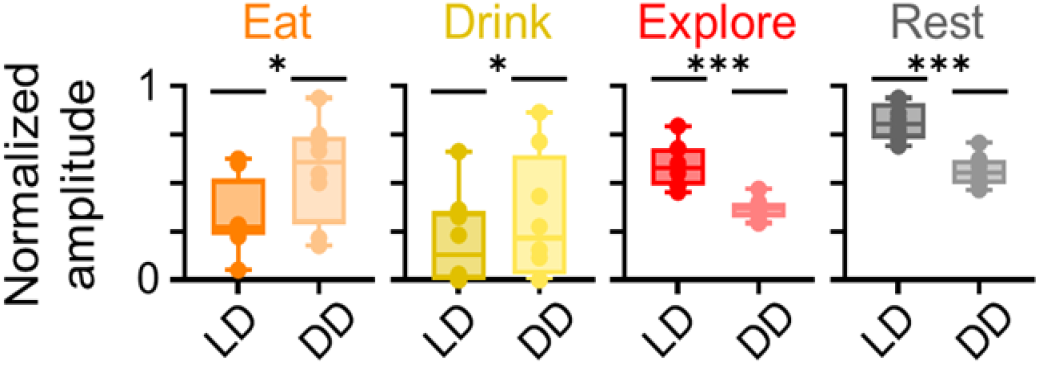
The amplitudes of exploratory and maintenance rhythms in wild-type male mice differ with ambient light. Rhythm amplitudes (in arbitrary units) for eating, drinking, exploring, and resting behaviors in individual wild-type male mice (n = 8) recorded in a 12 h:12 h light:dark cycle (LD) or in constant darkness (DD). Boxes indicate 25^th^ to 75^th^ percentiles. For visualization, rhythm amplitudes were normalized within each behavior. Statistical analysis performed on male and female wild-type and *Vip*-deficient mice in LD and in DD (see Supplementary Figure 3). Three-Way Repeated Measures ANOVA, *p < 0.05; *** p < 0.001.

**Figure 4.**
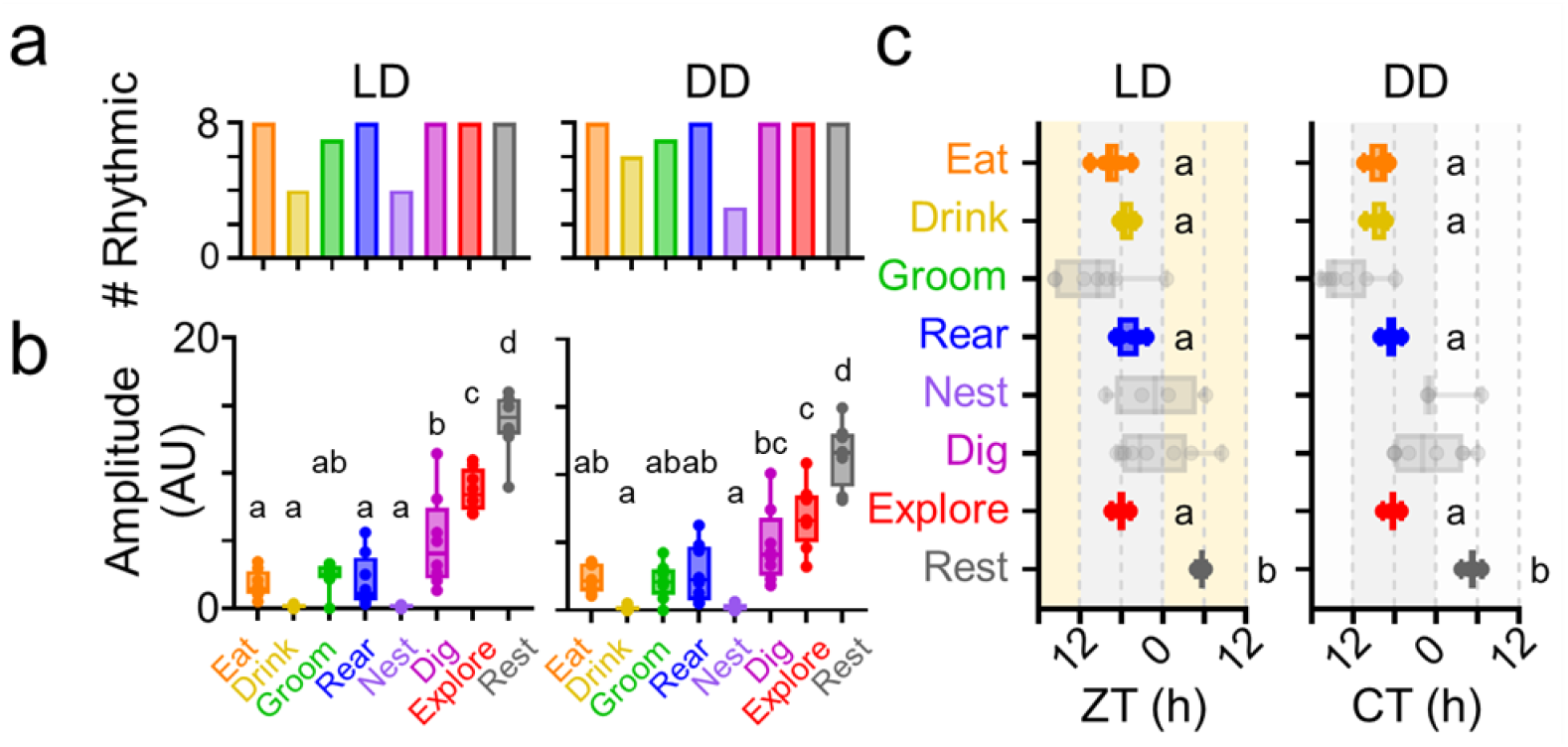
Wild-type female mice exhibit prominent diurnal and circadian rhythms in most behaviors. **a)** Number of individual wild-type female mice (n = 8) recorded in a 12 h:12 h lightdark cycle (LD) and in constant darkness (DD) scored as rhythmic for each behavior (ARSER, p < 0.05 or less). **b)** Rhythm amplitudes (in arbitrary units) for each behavior in LD and in DD. Boxes indicate 25^th^ to 75^th^ percentiles. Letters on each plot indicate similar amplitudes within each light cycle. (Two-Way Repeated Measures ANOVA, p < 0.05 or less). **c)** Peak times (in hours) for each behavior in individual mice in LD (yellow and dark gray background) and in DD (light and dark gray background). ZT. zeitgeber time. CT, circadian time. Boxes indicate 25^th^ to 75^th^ percentiles. Gray dots and boxes depict rhythms that did not have significant clustering of peak times across mice (Rayleigh test, p > 0.05). Letters on each plot indicate behaviors that peaked at similar times (Watson-Williams tests, p < 0.05 or less). Peak times depicted as linear for visualization.

### A subset of behavioral rhythms differs between male and female mice

We next wanted to determine if female mice exhibited behavioral rhythms that were similar to those of male mice and to identify which rhythms (if any) were sexually dimorphic. We found that, like wild-type male mice, most individual wild-type female mice exhibited prominent daily rhythms (ARSER, p < 0.05) in eating, drinking, grooming, rearing, nesting, digging, exploring, and resting behaviors that persisted in constant darkness (**Fig. 4a**). The amplitudes of exploring and resting rhythms were greater in LD (Three-Way Repeated Measures ANOVA, p < 0.001) and the amplitudes of eating and drinking rhythms were greater in DD (p < 0.05 or less; **Figs. 4b, 5a; Supplementary Fig. 3**). The phases of eating, drinking, rearing, exploring, and resting rhythms were highly similar across individual mice (Rayleigh test, p < 0.05) and did not differ between LD and DD (Multi-Way Circular ANOVA, p > 0.05; **Fig. 4c; Supplementary Fig. 4**). The phases and amplitudes of these rhythms did not differ significantly between male and female wild-type mice (p > 0.05).

Surprisingly, we found that nesting rhythms in individual wild-type female mice were much lower in amplitude than nesting rhythms in male mice in both LD and DD (Three-Way Repeated Measures ANOVA, p < 0.05; **Fig. 5b**). Conversely, digging rhythms in individual wild-type female mice had higher amplitudes than digging rhythms in male mice in both LD and DD (p < 0.05). The peak times of nesting, digging, and grooming rhythms in each female mouse were dispersed throughout the day and did not significantly cluster across mice (Rayleigh test, p > 0.05; **Fig. 4c; Supplementary Fig. 4**). However, on average, nesting and digging rhythms in individual female mice peaked later in the day than these rhythms in male mice (Multi-Way Circular ANOVA, p < 0.01; **Fig. 5c**). Together, these results demonstrate that while most behavioral rhythms are largely similar in male and female mice, there are clear sex differences in both nesting and digging rhythms.

**Figure 5.**
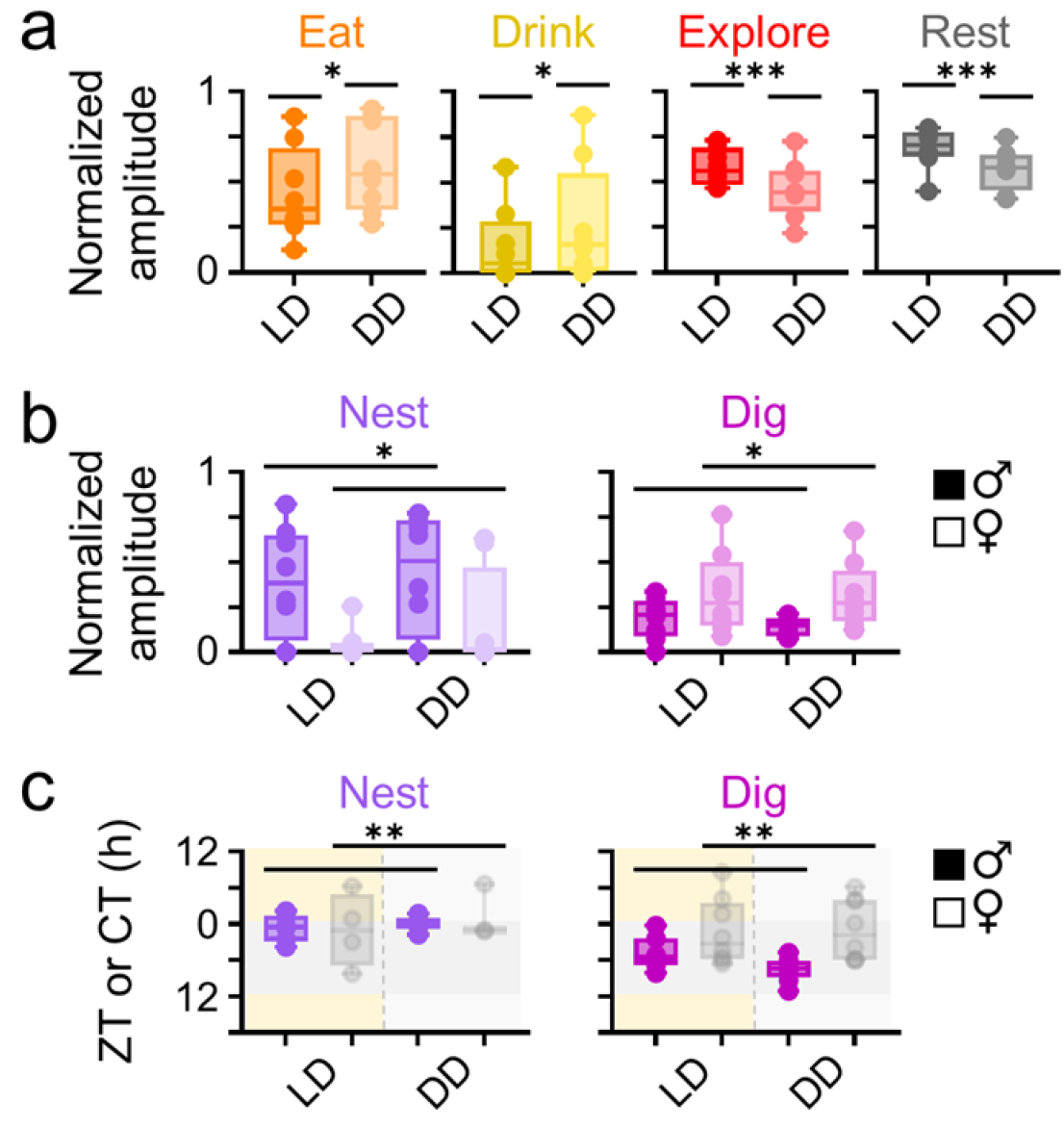
Nesting and digging rhythms differ between male and female mice. **a)** Rhythm amplitudes (in arbitrary units) for eating, drinking, exploring, and resting behaviors in individual wild-type female mice (n = 8) recorded in a 12 h:12 h light dark cycle (LD) or in constant darkness (DD). **b)** Rhythm amplitudes for nesting and digging behaviors in individual wild-type male (daik circles) and female (light circles) mice recorded in LD and in DD. **c)** Peak times (in hours) for nesting and digging behavior rhythms in individual wild-type male (dark circles) and female (light circles) mice recorded in LD (yellow and dark gray background) and in DD (light and dark gray background). Gray dots and boxes depict rhythms that did not have significant clustering of peak times across mice (Rayleigh test, p > 0.05). Boxes indicate 25^th^ to 75^th^ percentiles. For visualization, rhythm amplitudes were normalized within each behavior and peak times were depicted as linear. Statistical analysis performed on male and female wild-type and *Vip*-deficlent mice in LD and in DD (see Supplementary Figures 3 and 4). Three-Way Repeated Measures ANOVA and MultiWay Circular ANOVA, p < 0.05; **, p < 0.01*** p < 0.001.

### Diurnal rhythms in *Vip*^-/-^ mice are low amplitude and peak earlier in the day

It was immediately apparent from our ethograms that daily rhythms in most behaviors in *Vip*- deficient mice were markedly different from those in wild-type mice (**Fig. 1**). Surprisingly, we found that in LD, most individual male and female *Vip*^-/-^mice each exhibited significant rhythms (p < 0.05) in all assessed behaviors regardless of which rhythmicity detection method we used for analysis (**Figs. 6a, 7a; Supplementary Fig. 2**). We nevertheless determined that these diurnal rhythms in eating, drinking, rearing, exploring, and resting behaviors in individual *Vip*^-/-^mice were much lower in amplitude than these rhythms in wild-type mice (Three-Way Repeated Measures ANOVA, p < 0.05 or less; **Figs. 6b, 7b, 8a; Supplementary Fig. 3**). The phases of diurnal rhythms in most behaviors in both male and female *Vip*- /- mice were each highly similar across individual mice (Rayleigh test, p < 0.05; **Figs. 6c, 7c; Supplementary Fig. 4**). However, the peak times of diurnal rhythms in eating and grooming in male *Vip*^-/-^mice, and in drinking, grooming, rearing, and nesting in female *Vip*^-/-^mice, were each dispersed throughout the day and did not significantly cluster across mice (Rayleigh test, p > 0.05). In general, diurnal rhythms in eating, drinking, rearing, nesting, digging, exploring, and resting in individual *Vip*^-/-^mice peaked earlier in the day than these rhythms in wild-type mice (Multi-Way Circular ANOVA, p < 0.05 or less; **Fig. 8b**). Together, these results demonstrate that in a light:dark cycle, most behavioral rhythms in *Vip*-deficient mice are both lower in amplitude and phase advanced compared to those in wild-type mice.

**Figure 6.**
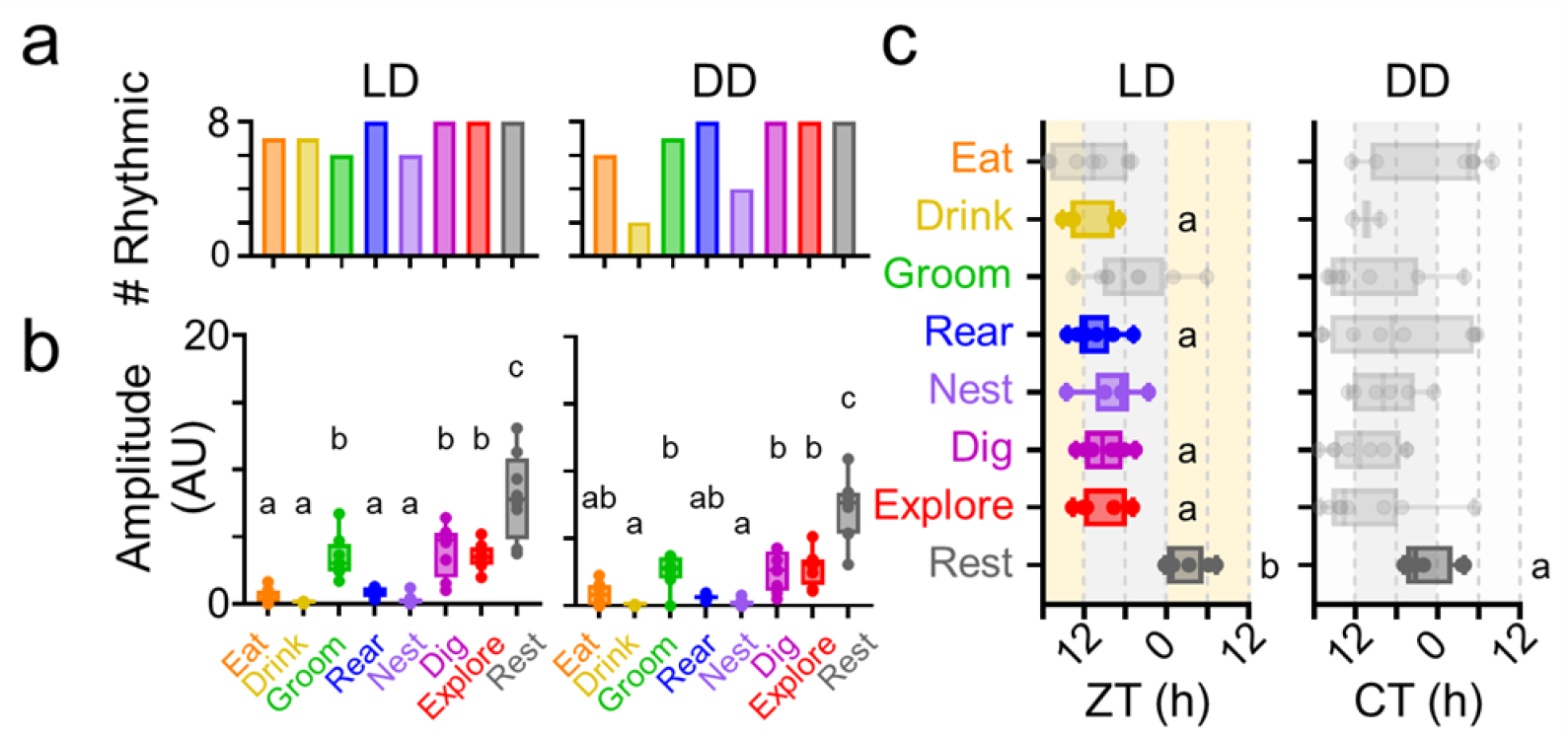
*Vip*-deficient male mice exhibit circadian rhythms in most behaviors that peak randomly throughout the day. **a)** Number of individual *Vip*-deficient male mice (n = 8) recorded in a 12 h: 12 h light dark cycle (LD) and in constant darkness (DD) scored as rhythmic for each behavior (ARSER, p < 0.05 or less). **b)** Rhythm amplitudes (in arbitrary units) for each behavior in LD and in DD. Boxes indicate 25^th^ to 75^th^ percentiles. Letters on each plot indicate similar amplitudes within each light cycle. (Two-Way Repeated Measures ANOVA, p < 0.05 or less). **c)** Peak times (in hours) for each behavior in individual mice in LD (yellow and dark gray background) and in DD (light and dark gray background). ZT, zeitgeber time. CT, circadian time. Boxes indicate 25^th^ to 75^th^ percentiles. Gray dots and boxes depict rhythms that did not have significant clustering of peak times across mice (Rayleigh test, p > 0.05). Letters on each plot indicate behaviors that peaked at similar times (Watson-Williams tests, p < 0.05 or less). Peak times depicted as linear for visualization.

**Figure 7.**
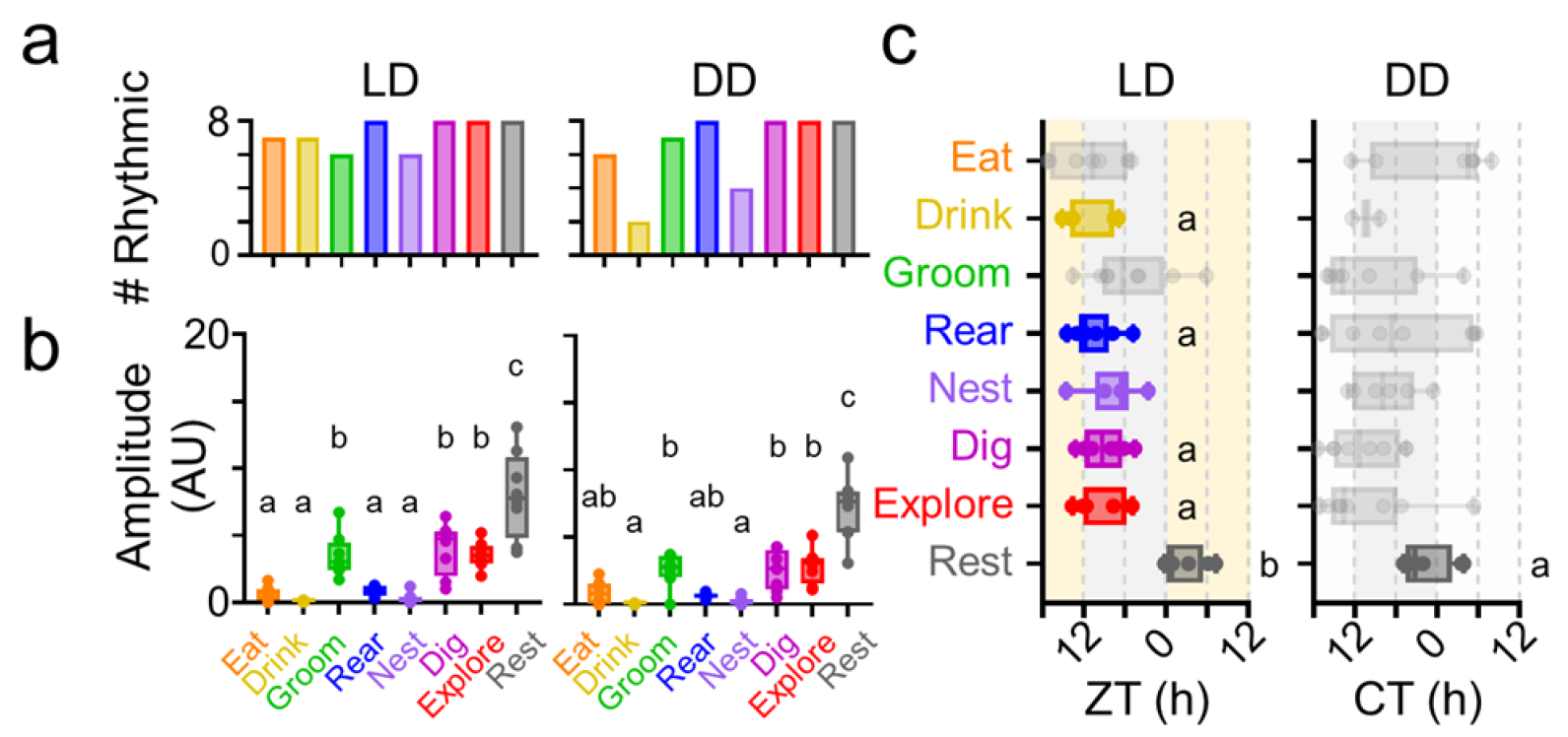
*Vip*-deficient female mice exhibit circadian rhythms in most behaviors that peak randomly throughout the day. **a)** Number of individual *Vip*-deficlent female mice (n = 8) recorded in a 12 h:12 h lightdark cycle (LD) and in constant darkness (DD) scored as rhythmic for each behavior (ARSER, p < 0.05 or less). **b)** Rhythm amplitudes (in arbitrary units) for each behavior in LD and in DD. Boxes indicate 25^th^ to 75^th^ percentiles. Letters on each plot indicate similar amplitudes within each light cycle (Two-Way Repeated Measures ANOVA, p < 0.05 or less). **c)** Peak times (in hours) for each behavior in individual mice in LD (yellow and dark gray background) and in DD (light and dark gray backgroun **d).** ZT, zeitgeber time. CT, circadian time. Boxes indicate 25^th^ to 75^th^ percentiles. Gray dots and boxes depict rhythms that did not have significant clustering of peak times across mice (Rayleigh test, p > 0.05). Letters on each plot indicate behaviors that peaked at similar times (Watson-Williams tests, p < 0.05 or less). Peak times depicted as linear for visualization.

**Figure 8.**
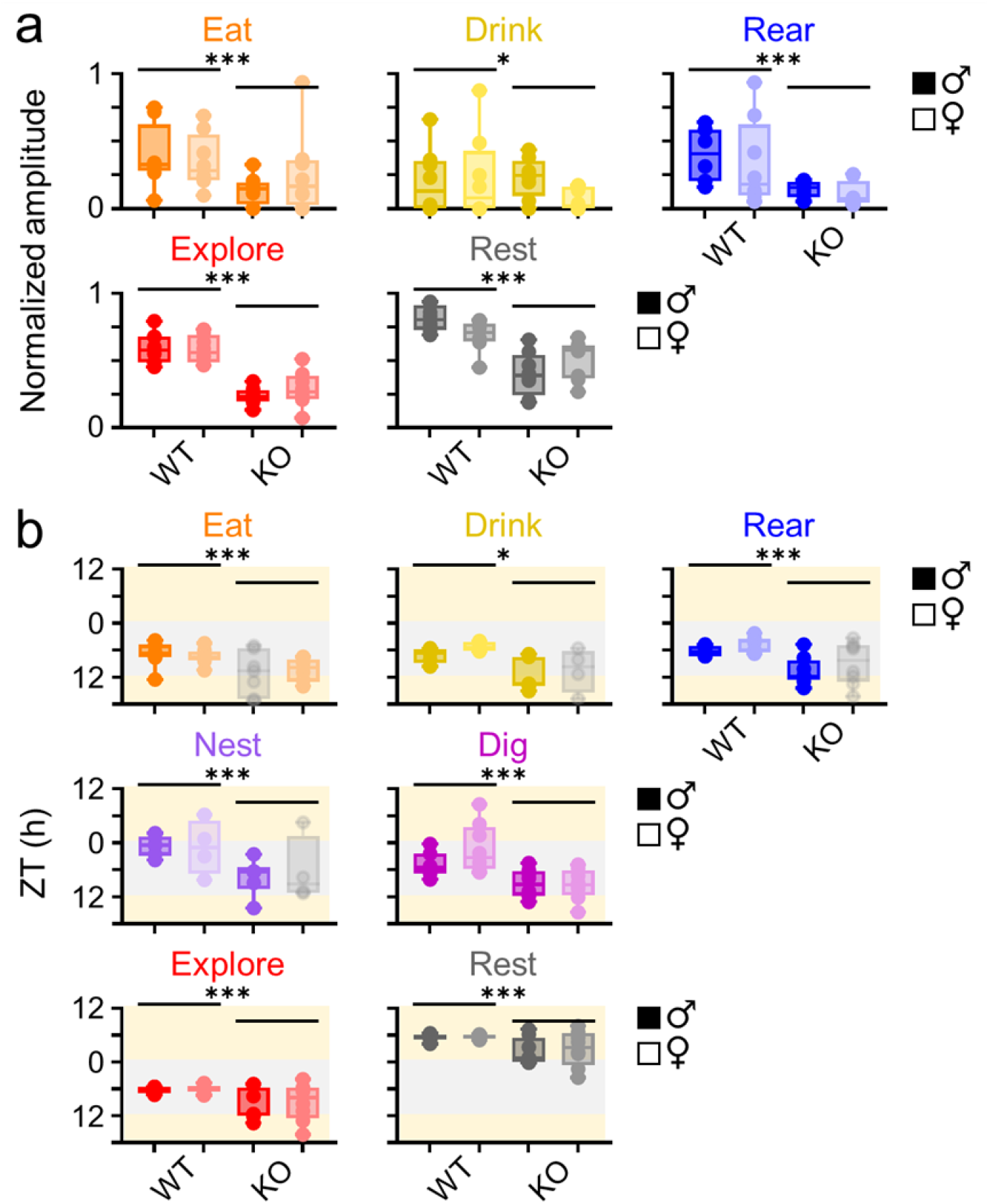
*Vip*-deficient mice exhibit low amplitude, phase-advanced rhythms in LD. **a)** Rhythm amplitudes (in arbitrary units) for eating, drinking, rearing, exploring, and resting behaviors in individual wild-type and *Vip*-deficient male (dark circles) and female (light circles) mice (n = 8 mice per condition) recorded in a 12 h:12 h lightdark cycle (LD). **b)** Peak times (in hours) for eating, drinking, rearing, nesting, digging, exploring, and resting behavior rhythms in individual wild-type and *Vip*-deficient male (dark circles) and female (light circles) mice recorded in LD (yellow and dark gray background). Gray dots and boxes depict rhythms that did not have significant clustering of peak times across mice (Rayleigh test, p > 0.05). Boxes indicate 25^th^ to 75^th^ percentiles. For visualization, rhythm amplitudes were normalized within each behavior and peak times were depicted as linear. Statistical analysis performed on male and female wild-type and Vfp-deficient mice in LD and in DD (see Supplementary Figures 3 and 4). Three-Way Repeated Measures ANOVA and Multi-Way Circular ANOVA, p < 0.05 *** p < 0.001.

### Circadian rhythms in *Vip*^-/-^mice peak randomly throughout the day

We found that, just as we observed in LD, most individual male and female *Vip*^-/-^mice in DD each exhibited significant rhythms (p < 0.05) in all assessed behaviors regardless of which rhythmicity detection method we used for analysis (**Figs. 6a, 7a; Supplementary Fig. 2**). Drinking rhythms were absent in DD in most individual *Vip*^-/-^mice. As with wild-type mice, the amplitudes of exploring and resting rhythms in individual *Vip*^-/-^mice were greater in LD and the amplitude of eating rhythms was greater in DD (Three-Way Repeated Measures ANOVA, p < 0.05 or less; **Fig. 9a; Supplementary Fig. 3**). We also found that, just as we observed in LD, circadian rhythms in eating, drinking, rearing, exploring, and resting behaviors in individual *Vip*^-/-^mice were again much lower in amplitude than these rhythms in wild-type mice (Three-Way Repeated Measures ANOVA, p < 0.05 or less; **Fig. 9b**). However, critically, the peak times of circadian rhythms in all behaviors but resting in male *Vip*^-/-^mice, and in all behaviors in female *Vip*^-/-^mice, were each dispersed throughout the day and did not significantly cluster across mice (Rayleigh test, p > 0.05; **Figs. 6c, 7c; Supplementary Fig. 4**). Circadian rhythms in resting behavior in individual *Vip*^-/-^male mice had similar phases (p < 0.05) and peaked earlier in the day than these rhythms in wild-type mice (Multi-Way Circular ANOVA, p < 0.001; **Fig. 9c**). Together, these results demonstrate that in constant darkness, behavioral rhythms in *Vip*-deficient mice peak randomly throughout the day.

**Figure 9.**
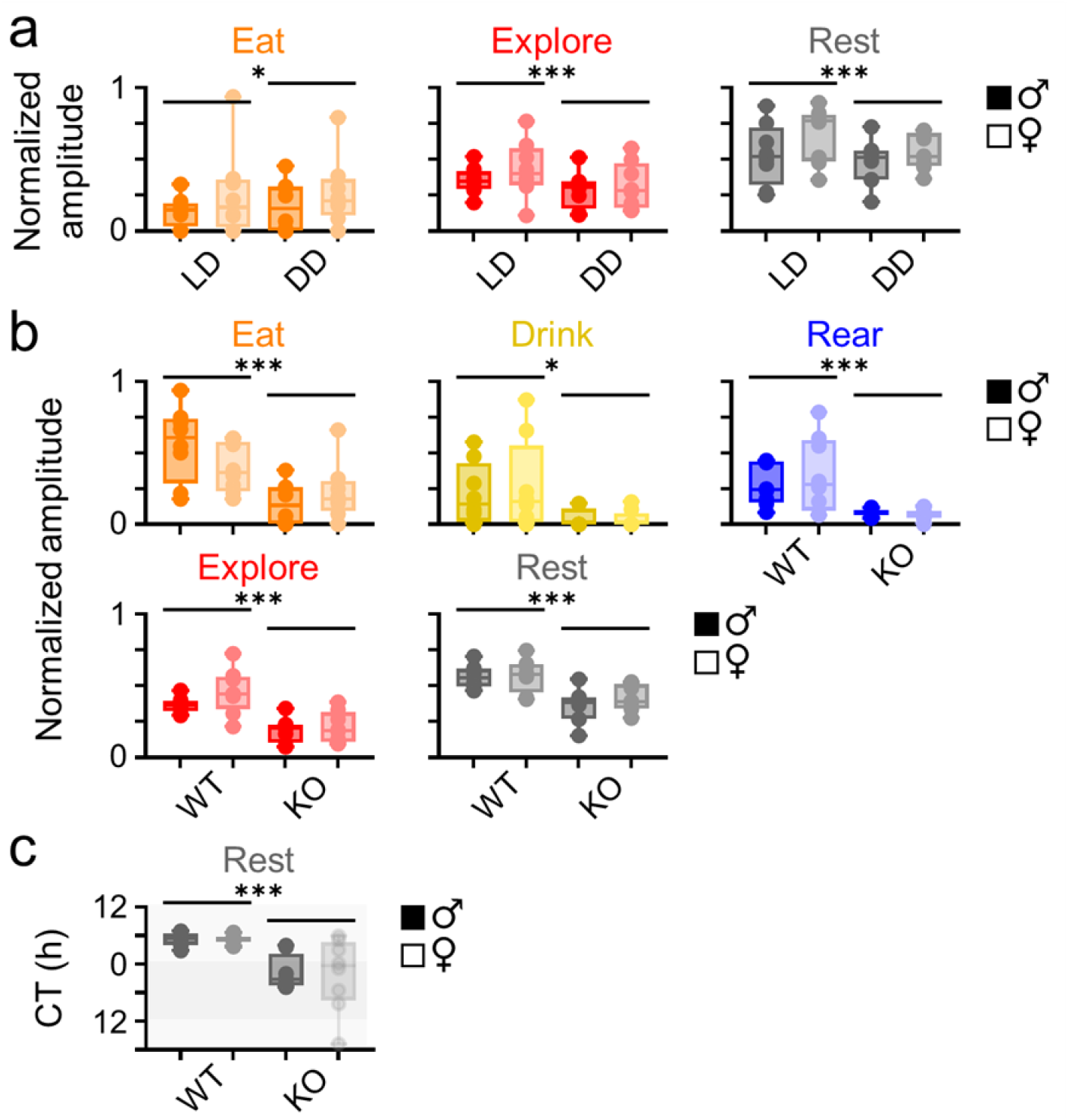
*Vip*-deficient mice exhibit low amplitude rhythms in DD. **a)** Rhythm amplitudes (in arbitrary units) for eating, exploring, and resting behaviors in individual *Vip*-deficient male (dark circles) and female (light circles) mice (n = 8 per condition) recorded in a 12 h: 12 h light:dark cycle (LD) or in constant darkness (DD). **b)** Rhythm amplitudes (in arbitrary units) for eating, drinking, rearing, exploring, and resting behaviors in Individual wild-type and *Vip*-deficient male (dark circles) and female (light circles) mice (n = 8 mice per condition) recorded in DD. **b)** Peak times (in hours) for resting behavior rhythms in individual wild-type and *Vip*-deficient male (dark circles) and female (light circles) mice recorded in DD (light and dark gray background). Gray dots and boxes depict rhythms that did not have significant clustering of peak times across mice (Rayleigh test, p > 0.05). Boxes indicate 25^th^ to 75^th^ percentiles. For visualization, rhythm amplitudes were normalized within each behavior and peak times were depicted as linear. Statistical analysis performed on male and female wild-type and *Vip*-deficient mice in LD and in DD (see Supplementary Figures 3 and 4). Three-Way Repeated Measures ANOVA and Multi-Way Circular ANOVA, p < 0.05; *** p < 0.001.

### Time budgets and transition probabilities differ with VIP signaling, by sex, and by light cycle

The high temporal resolution of our data allowed us to determine the temporal sequence of behaviors on the seconds-to-minutes timescale over multiple days. We therefore calculated time budgets (the total time a mouse spent performing a given behavior across 48 h; **Supplementary Fig. 5a**) and transition probabilities (the probability that a given behavior would follow an “initial behavior” within 10 seconds; **Supplementary Fig. 6a**) for each behavior (Lehner, 1998). We found that for most behaviors, time budgets did not differ by genotype (3/8 behaviors differed between wild-type and *Vip*^-/-^mice; Three-Way Repeated Measures ANOVA, p < 0.05 or less) or sex (2/8 behaviors differed between male and female mice; p < 0.05 or less; **Supplementary Fig. 5b**). However, time budgets strongly differed by light cycle (6/8 behaviors differed between LD and DD, p < 0.05 or less). These differences were largely driven by the light cycle instead of the day of recording *per se*, as only exploring behavior showed a significant difference in time budget between days in LD (p < 0.001) or DD. Conversely, we found that for most behaviors, transition probabilities did not differ by light cycle (3/8 behaviors differed between LD and DD; Three-Way Repeated Measures ANOVA, p < 0.05 or less; **Supplementary Fig. 6b**). However, transition probabilities for multiple behaviors differed by genotype (5/8 behaviors differed between wild-type and *Vip*^-/-^mice; p < 0.05 or less) and sex (6/8 behaviors differed between male and female mice; p < 0.05 or less). Together, these results demonstrate that time budgets are most strongly influenced by the light cycle in which the mice were recorded, but the probability of an animal transitioning from one behavior to another is most strongly influenced by VIP signaling and sex.

## Discussion

To test the hypothesis that VIP signaling, sex, and ambient light affect daily patterns of behavior, we used machine learning to measure the temporal distribution of multiple behaviors in male and female wild type and *Vip*^-/-^mice over several days in LD and in DD. We conclude that circadian rhythms in both established and novel behaviors differ with and without VIP signaling, between males and females, and in a light cycle or in constant darkness. Our identification of these novel behavioral rhythms and our implementation of a method to automatically classify daily rhythms in behavior will provide a foundation for future studies investigating the temporal organization of other complex behaviors.

We found that each measured behavior was typically rhythmic in individual animals of either genotype or sex in both LD and in DD regardless of which rhythmicity analysis method we used (Refinetti et al., 2007; Zielinski et al., 2014; Wu et al., 2016). These algorithms may have a propensity for false positives with our data because, for the large number of time points we recorded within a limited sampling window (345,600 frames over 96 h), periods of inactivity and activity could inadvertently make a behavior appear rhythmic. However, the rhythmicity we observed in a majority of behaviors in a majority of individual mice is consistent with several previous studies. For example, wheel-running activity is rhythmic in most *Vip*^-/-^mice in LD and for the first several days in DD (Aton et al., 2005). It is possible that the rhythms we observed in some or all behaviors in our *Vip*^-/-^mice would dampen over time in DD.

We found that several behavioral rhythms differed between wild-type and *Vip*-deficient animals. For instance, rhythms in eating, drinking, rearing, exploring, and resting behaviors were each dramatically lower in amplitude in our *Vip*^-/-^mice than in our wild-type mice. This is consistent with previous studies that found attenuated wheel-running activity, sleep, and feeding rhythms in *Vip*^-/-^mice, but alterations in drinking and rearing rhythm amplitudes in these animals have not been previously reported (Colwell et al., 2003; Bechtold et al., 2008; Hu et al., 2011). Similarly, rhythms in eating, drinking, rearing, nesting, exploring, and resting behaviors peaked earlier in the day in our *Vip*^-/-^mice than in our wild-type mice. This is again consistent with previous studies that found that *Vip*^-/-^mice have phase-advanced wheel-running activity and sleep rhythms, but phase differences in eating, drinking, rearing, and nesting have not been previously reported (Colwell et al., 2003; Hu et al., 2011). Importantly, this study is the first to simultaneously measure each of these behaviors in *Vip*-deficient mice. This parallel analysis is critical to understand how disrupted VIP signaling in the SCN affects the temporal sequencing of multiple behaviors.

We also found that rhythms in digging and nesting behavior differed between male and female mice. The sex differences we observed in the amplitude and phase of these behavioral rhythms are entirely novel and have not been previously reported. However, other studies have found that female mice show a longer duration of wheel-running activity (comparable to our “exploring” behavior) than male mice in DD, but male mice show a greater precision of wheel-running activity than female mice in LD (Kuljis et al., 2013). We were unable to directly compare our results to these findings because we only measured behaviors over two days in LD and in DD. However, we did observe that the duration of exploring rhythms in female mice was indeed slightly longer than in male mice (14.6 ± 0.5 h vs. 13.7 ± 0.6 h; unpaired Welch’s *t* test, p < 0.01). These results emphasize that measuring circadian rhythms in behaviors other than locomotor activity can reveal critical unseen sex differences in behavior.

Intriguingly, the VIP-, sex, and light-dependent differences we observed were only present in a subset of a mouse’s entire behavioral repertoire. For instance, we only observed a VIP signaling-dependent difference in the amplitude of daily rhythms in five out of eight measured behaviors and a sex difference in the amplitude and phase of daily rhythms in two out of eight measured behaviors. This suggests that the brain regions and neural circuits that regulate each of these behaviors may be differentially influenced by the daily timing signal that originates from the SCN. For example, the circuit that regulates digging rhythms may be relatively robust to a desynchronized input arising from a *Vip*-deficient SCN, but the circuit that regulates exploring rhythms may be more directly affected by *Vip* deficiency. Future experiments should investigate the role of local clocks in brain circuits that are known to be associated with each of these behaviors.

We found that VIP signaling, sex, and ambient light also affected how the similar the phases of behavioral rhythms were across mice. For instance, the phases of most rhythms were highly similar across individual *Vip*^-/-^mice in LD but were dispersed throughout the day in DD. This suggests that *Vip*-deficient mice fail to entrain to the LD cycle, which causes their behavioral rhythms to peak at random times in DD. We also found that the phases of nesting and digging rhythms were highly similar across individual male mice but were dispersed throughout the day in individual female mice. Indeed, nesting rhythms in male mice had their own unique phase compared to all other behaviors. The mechanism underlying this phase dispersal across individual female, but not male mice in some, but not all behavioral rhythms is unclear, but may be due to changes in specific circuits that respond differently to circulating sex hormones.

We also identified VIP signaling-, sex-, and light-dependent differences in time budgets and transition probabilities for several behaviors. For instance, we observed that mice switched their time budget distribution from exploratory behaviors in LD to maintenance (eating and drinking) behaviors in DD. Similarly, we observed that the time budgets for grooming and digging behaviors differed between male and female mice. This is consistent with previous studies that identified sex differences in these behaviors within a much shorter temporal window (Geuther et al., 2021; Pond et al., 2021). Surprisingly, we also observed differences in time budgets and transition probabilities for several behaviors between *Vip*-deficient and wild-type mice. Time budgets and transition probabilities are each ostensibly independent of circadian time. Consequently, SCN desynchrony caused by disrupted VIP signaling should theoretically have no effect on these measurements. There is limited evidence that the SCN can influence certain behaviors independently of its role in rhythm generation (Yu et al., 2017). Alternatively, VIP is also expressed in other neurons, such as those in the olfactory bulb and cortex (Lein et al., 2007). Disrupted VIP signaling in these circuits could result in our observed differences in behavior.

In this study, we used machine learning to automatically identify differences in the temporal organization of behavior due to VIP signaling, sex, and ambient light. This approach can be readily expanded to address other critical questions in neuroscience and circadian biology, including the ethological investigation of other behavioral rhythms in videos of mice recorded in the laboratory and, potentially, in the wild. Notably, machine learning can also be used for the rapid circadian phenotyping of mice with different genotypes or disorders. Current approaches almost universally measure changes to wheel-running activity rhythms as evidence that a mutation, gene, or drug influences circadian behavior. Here, we found that some, but not all, behavioral rhythms differ by sex and with VIP signaling. It is therefore likely that a given experimental treatment may cause circadian alterations in behaviors other than, or in addition to, wheel-running activity. Machine learning can be used to study these circadian behaviors that were previously difficult or impossible to observe.

## Methods

### Animals

Prior to recording, we group-housed male and female wild-type (*Vip*^+/+^, n = 8 per sex) and *Vip*-deficient (*Vip*^-/-^, n = 8 per sex) mice (Colwell et al., 2003) in their home cages in a 12 h:12 h light:dark cycle (LD, where lights on is defined as zeitgeber time (ZT) 0; light intensity ∼2 x 10^14^ photons/cm^2^/s) at constant temperature (∼23°C) and humidity (∼40%) with food and water provided ad libitum. All mice were between 6 and 12 weeks old at the time of recording. All experiments were approved by and performed in accordance with the guidelines of Texas A&M University and Washington University’s Institutional Animal Care and Use Committees.

### Experimental housing

We transferred individual mice from their home cages to custom-built recording cages inside light-tight, temperature-and-humidity controlled circadian cabinets for the duration of our experiments. The cages are built out of transparent 6 mm-wide acrylic and are approximately 21 cm in length, width, and height. In one of the cage walls, we installed an acrylic food hopper with a slotted opening of around 15 x 10 cm that we filled with food pellets. On the outside of the wall immediately adjacent to the food hopper we installed a water bottle with a metal spout that protruded about 1.5 cm into the cage. We covered the cage floor (around 440 cm^2^ of explorable space) with about 1 cm of standard corn cob bedding and placed half of a cotton nestlet in the center of the cage. In the light phase in LD, we illuminated the cabinets with broad-spectrum white light (∼6 x 10^13^ photons/cm^2^/s measured at the cage floor). In the dark phase in LD and in constant darkness (DD), we illuminated the cabinets with dim red light (660 nm, ∼3 x 10^13^ photons/cm^2^/s). We acclimated mice in the recording cages for 20-24 hours before the start of each experiment.

### Video recording

We positioned networked video cameras (DCS-932L and DCS-933L, D-Link) equipped with fish-eye lenses directly above the recording cages such that all four corners of the cage, the food hopper, and water spout were each visible in the recorded video and the mouse and nesting material were in focus (**Supplementary Fig. 1a**). We continually recorded grayscale videos (640 x 480 pixel resolution, 30 frames per second) of singly-housed mice as they freely behaved continually over four days, two days in 12h:12h LD and two days in DD. We then exported the recorded videos as .asf files using D-ViewCam software, downsampled the videos to 1 fps and converted the videos to .mp4 files using the open-source video editing program FFMpeg. Finally, we cropped the converted videos to the extent of the cage floor using the open-source video editing program Handbrake.

### Model training and inference

We installed Deep Ethogram from source on a custom-built machine learning computer (12-core AMD Ryzen 9 5900X CPU, 32 GB RAM, NVIDIA GeForce RTX 3090 with 24 GB VRAM) according to the DeepEthogram Github installation guide (http://github.com/jbohnslav/deepethogram/) (Bohnslav et al., 2021). To train the DeepEthogram models, we first chose eight complex motor behaviors that encompass the majority of a mouse’s daily activity: eating, drinking, grooming, rearing, nesting, digging, exploring, and resting. We manually labeled each frame of the training videos (three 24 h, 86,400 frame-long videos of wild-type male mice and five 10 min, 600 frame-long videos of wild-type female and male and female *Vip*^-/-^mice) using the GUI based on consensus between three trained classifiers and the behavioral descriptions listed on the Stanford University Mouse Ethogram index (Garner, 2017). To create our machine-learning model, we first trained the “flow generator” convolutional neural network (CNN), which uses the MotionNet architecture to estimate motion (“optic flow”) from our training video frames. The flow generator CNN is pre-trained on the Kinetics700 video dataset and requires no user input. Next, we trained a two-stream CNN, a “feature extractor,” on the output from the flow generator and our manually-labeled frames. The feature extractor, also pre-trained on the Kinetics700 video dataset, uses the ResNet50 architecture to determine the probability of a behavior being present in a particular frame based on a low-dimensional set of temporal (optic flow) and spatial (labeled frames) features. Finally, we used our feature extractor outputs to train a “sequence model” CNN that uses the Temporal Gaussian Mixture CNN which has a large temporal receptive field and can thus further refine the model predictions using a longer timescale to provide “context” for a given behavior frame. To prevent the model from predicting multiple behaviors for a given frame, we changed the final activation value for our feature extractor and sequence model CNNs from “sigmoid” to “softmax” during training. A softmax, or normalized exponential activation, function requires that the probability of predicted behaviors on a given frame sum to one and thus, for our inferred videos, each frame was uniquely labeled as one behavior. The trained model was then used to predict behaviors from our experimental video recordings (**Supplementary Fig. 1a**).

### Analysis

We determined circadian rhythmicity using three methods (**Supplementary Fig. 3**): Cosinor analysis in Matlab (Refinetti et al., 2007)and Lomb-Scargle Periodogram and ARSER in Metacycle (Wu et al., 2016). We also performed empirical JTK_Cycle in BioDare2 (Zielinski et al., 2014) on data in 5 min bins instead of on the entire dataset because the current release of JTK_Cycle struggles to handle data of this length. We ultimately used rhythmicity, amplitude, and phase predictions by ARSER to perform comparisons of circadian rhythms. We calculated rhythm amplitudes and peak times for individual mice using the amplitude and phase values determined by ARSER. We calculated total activity by summing the total frames exhibiting a given behavior for individual mice in LD and in DD. We calculated transition scores, which we defined as the probability that a given behavior will follow another behavior within 10 or fewer seconds, using a custom MATLAB script.

We performed the following statistical tests in Prism 9.0 (GraphPad, San Diego, CA): Two-Way ANOVA, Two-Way Repeated Measures ANOVA, Three-Way Repeated Measures ANOVA, post-hoc Sidak’s multiple comparisons test. We performed the following statistical analyses using the Circular Statistics Toolbox in Matlab (Berens, 2009) (Mathworks, Natick, MA): Rayleigh test, multi-way circular ANOVA, and Watson-Williams test. We performed the following statistical analysis using the Statistics Toolbox in Matlab: multi-way ANOVA with post hoc Tukey’s HSD test. We used Shapiro–Wilk and Brown– Forsythe tests to test for normality and equal variances, defined α as 0.05, and presented all data as mean ± SEM.

## References

Ancoli-Israel S, Cole R, Alessi C, Chambers M, Moorcroft W, Pollak CP (2003) The role of actigraphy in the study of sleep and circadian rhythms. Sleep 26:342–392.

Aton SJ, Colwell CS, Harmar AJ, Waschek J, Herzog ED (2005) Vasoactive intestinal polypeptide mediates circadian rhythmicity and synchrony in mammalian clock neurons. Nat Neurosci 8:476–483.

Bechtold DA, Brown TM, Luckman SM, Piggins HD (2008) Metabolic rhythm abnormalities in mice lacking VIP-VPAC2 signaling. Am J Physiol Regul Integr Comp Physiol 294:R344– R351.

Berens P (2009) CircStat: a MATLAB toolbox for circular statistics. J Stat Softw 31:1–21.

Bohnslav JP, Wimalasena NK, Clausing KJ, Dai YY, Yarmolinsky DA, Cruz T, Kashlan AD, Chiappe ME, Orefice LL, Woolf CJ, Harvey CD (2021) DeepEthogram, a machine learning pipeline for supervised behavior classification from raw pixels. Elife 10 63377.

Colwell CS, Michel S, Itri J, Rodriguez W, Tam J, Lelievre V, Hu Z, Liu X, Waschek JA (2003) Disrupted circadian rhythms in VIP- and PHI-deficient mice. Am J Physiol Regul Integr Comp Physiol 285:R939–R949.

Garner J (2017) Mouse Ethogram. Available at: https://mousebehavior.org/ethogram/ [Accessed August 17, 2022].

Geuther BQ, Peer A, He H, Sabnis G, Philip VM, Kumar V (2021) Action detection using a neural network elucidates the genetics of mouse grooming behavior. Elife 10 63207.

Hastings MH, Maywood ES, Brancaccio M (2018) Generation of circadian rhythms in the suprachiasmatic nucleus. Nat Rev Neurosci 19:453–469.

Hu W-P, Li J-D, Colwell CS, Zhou Q-Y (2011) Decreased REM sleep and altered circadian sleep regulation in mice lacking vasoactive intestinal polypeptide. Sleep 34:49–56.

Jones JR, Simon T, Lones L, Herzog ED (2018) SCN VIP Neurons Are Essential for Normal Light-Mediated Resetting of the Circadian System. J Neurosci 38:7986–7995.

Joye DAM, Evans JA (2022) Sex differences in daily timekeeping and circadian clock circuits. Semin Cell Dev Biol 126:45–55.

Jud C, Schmutz I, Hampp G, Oster H, Albrecht U (2005) A guideline for analyzing circadian wheel-running behavior in rodents under different lighting conditions. Biol Proced Online 7:101–116.

Krizo JA, Mintz EM (2014) Sex differences in behavioral circadian rhythms in laboratory rodents. Front Endocrinol 5:234.

Kuljis DA, Gad L, Loh DH, MacDowell Kaswan Z, Hitchcock ON, Ghiani CA, Colwell CS (2016) Sex Differences in Circadian Dysfunction in the BACHD Mouse Model of Huntington’s Disease. PLoS One 11:e0147583.

Kuljis DA, Loh DH, Truong D, Vosko AM, Ong ML, McClusky R, Arnold AP, Colwell CS (2013) Gonadal- and sex-chromosome-dependent sex differences in the circadian system. Endocrinology 154:1501–1512.

Lee TM, Hummer DL, Jechura TJ, Mahoney MM (2004) Pubertal development of sex differences in circadian function: an animal model. Ann N Y Acad Sci 1021:262–275.

Lehner PN (1998) Handbook of Ethological Methods. Cambridge University Press.

Lein ES et al. (2007) Genome-wide atlas of gene expression in the adult mouse brain. Nature 445:168–176.

Loh DH, Abad C, Colwell CS, Waschek JA (2008) Vasoactive intestinal peptide is critical for circadian regulation of glucocorticoids. Neuroendocrinology 88:246–255.

Mazuski C, Abel JH, Chen SP, Hermanstyne TO, Jones JR, Simon Doyle FJ 3rd, Herzog ED (2018) Entrainment of Circadian Rhythms Depends on Firing Rates and Neuropeptide Release of VIP SCN Neurons. Neuron 99:555–563.e5.

Paul S, Hanna L, Harding C, Hayter EA, Walmsley L, Bechtold DA, Brown TM (2020) Output from VIP cells of the mammalian central clock regulates daily physiological rhythms. Nat Commun 11:1–14.

Pendergast JS, Branecky KL, Yang W, Ellacott KLJ, Niswender KD, Yamazaki S (2013) High-fat diet acutely affects circadian organisation and eating behavior. Eur J Neurosci 37:1350–1356.

Pond HL, Heller AT, Gural BM, McKissick OP, Wilkinson MK, Manzini MC (2021) Digging behavior discrimination test to probe burrowing and exploratory digging in male and female mice. J Neurosci Res 99:2046–2058.

Refinetti R, Lissen GC, Halberg F (2007) Procedures for numerical analysis of circadian rhythms. Biol Rhythm Res 38:275–325.

Schroeder A, Loh DH, Jordan MC, Roos KP, Colwell CS (2011) Circadian regulation of cardiovascular function: a role for vasoactive intestinal peptide. Am J Physiol Heart Circ Physiol 300:H241–H250.

Schwartz WJ, Zimmerman P (1990) Circadian timekeeping in BALB/c and C57BL/6 inbred mouse strains. J Neurosci 10:3685– 3694.

Todd WD, Venner A, Anaclet C, Broadhurst RY, De Luca R, Bandaru SS, Issokson L, Hablitz LM, Cravetchi O, Arrigoni E, Campbell JN, Allen CN, Olson DP, Fuller PM (2020) Suprachiasmatic VIP neurons are required for normal circadian rhythmicity and comprised of molecularly distinct subpopulations. Nat Commun 11:4410.

Walton JC, Bumgarner JR, Nelson RJ (2022) Sex differences in circadian rhythms. Cold Spring Harb Perspect Biol 14.

Wu G, Anafi RC, Hughes ME, Kornacker K, Hogenesch JB (2016) MetaCycle: an integrated R package to evaluate periodicity in large scale data. Bioinformatics 32:3351–3353.

Yerushalmi S, Green RM (2009) Evidence for the adaptive significance of circadian rhythms. Ecol Lett 12:970–981.

Yu Y-Q, Barry DM, Hao Y, Liu X-T, Chen Z-F (2017) Molecular and neural basis of contagious itch behavior in mice. Science 355:1072–1076.

Zhang C, Li H, Han R (2020) An open-source video tracking system for mouse locomotor activity analysis. BMC Res Notes 13:48.

Zielinski T, Moore AM, Troup E, Halliday KJ, Millar AJ (2014) Strengths and limitations of period estimation methods for circadian data. PLoS One 9:e96462.

