## Supplementary Figures for "Circadian rhythms in multiple behaviors depend on sex, neuropeptide signaling, and ambient light"

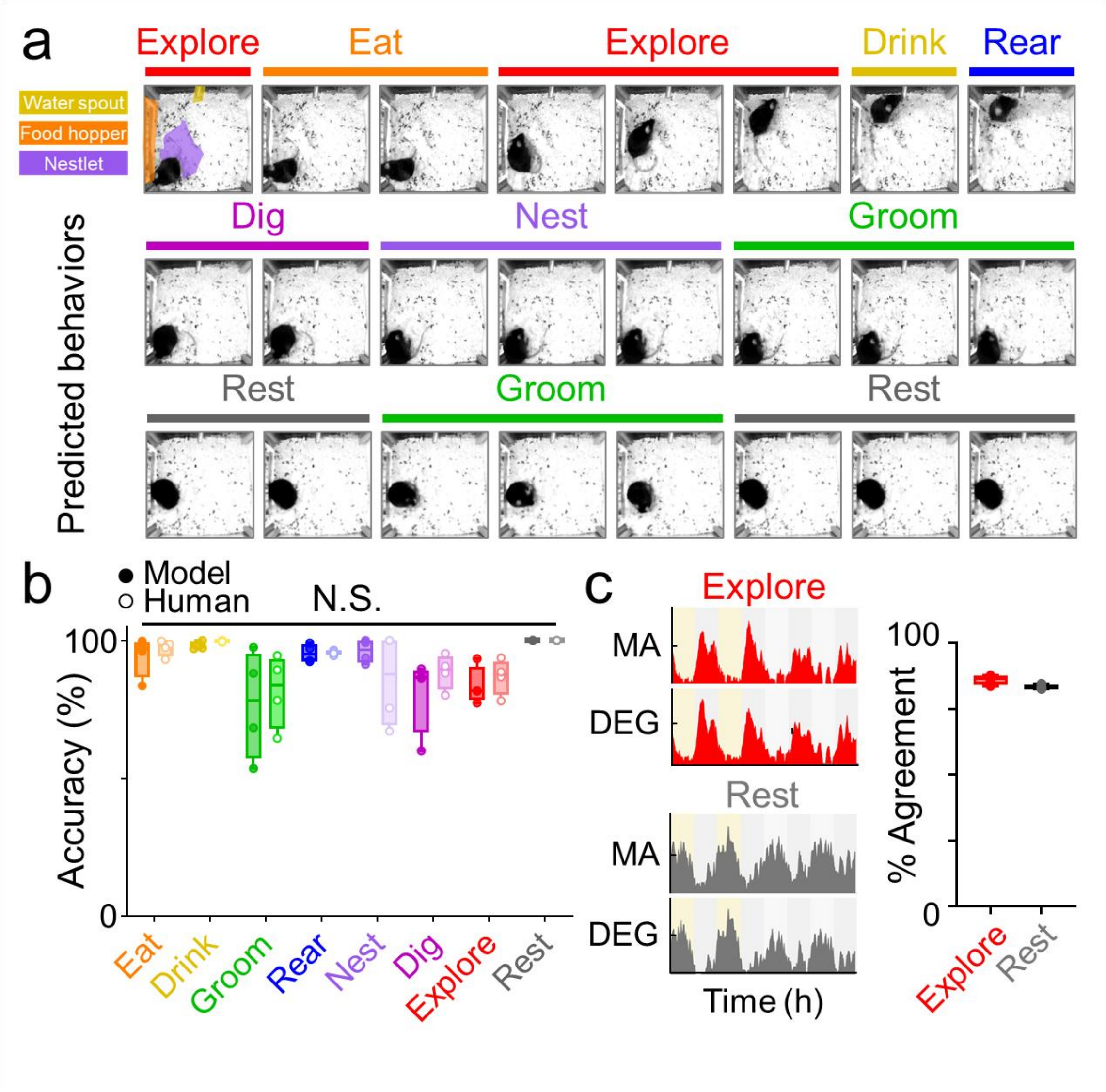


**Supplementary Figure 1. Validation of automated machine learning classification of complex behaviors**. **a)** Model-predicted labels for three 8-second behavior bouts (top to bottom). Bottom bout was recorded during the light phase; top and middle bouts were recorded during the night phase under dim red light. **b)** For each behavior, the percentage of frames from each of four videos labeled by our model (solid circles) or human classifiers (open circles) that were identical to the reference standard in each video. There was no significant difference between our model's predictions and behavior labels by human classifiers (Two-Way ANOVA,

$p \geq 0.500$ or greater for each behavior). **c)** *Left*, representative 96 h actograms for “exploring” and “resting” behaviors predicted by our model (DEG) and an independent automated behavior analysis program (MouseActivity, MA). *Right*, percentage of frames from each of four videos that were labeled identically as “exploring” or “resting” by our model or the MouseActivity program.

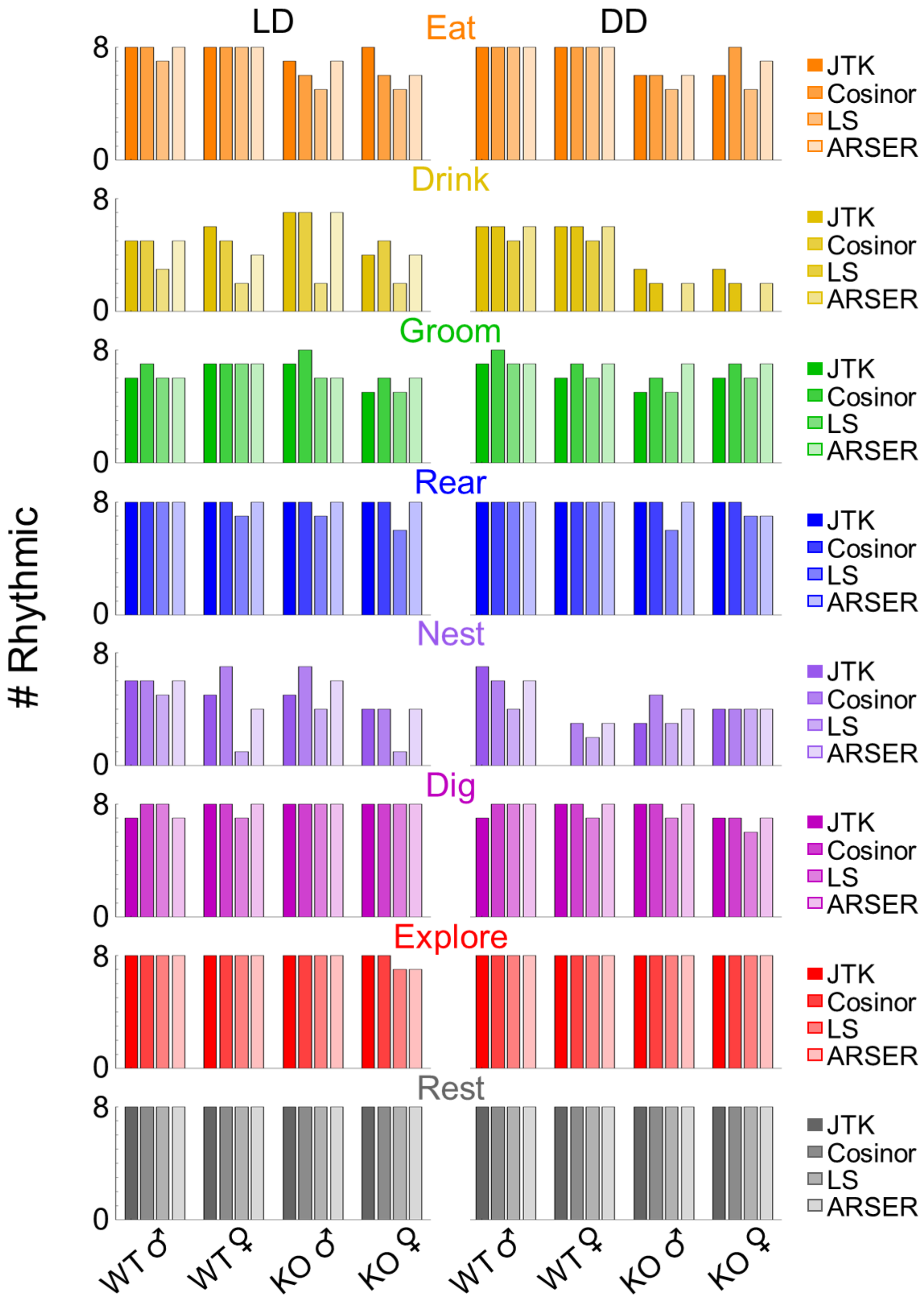


**Supplementary Figure 2. Comparison of different rhythmicity analyses.** Fraction of individual wild-type and *Vip*$^{-/-}$ male and female mice in a 12 h:12 h light:dark cycle (LD) and in constant darkness (DD) scored as rhythmic for each behavior using empirical JTK Cycle, Cosinor, Lomb-Scargle Periodogram, or ARSER analyses.

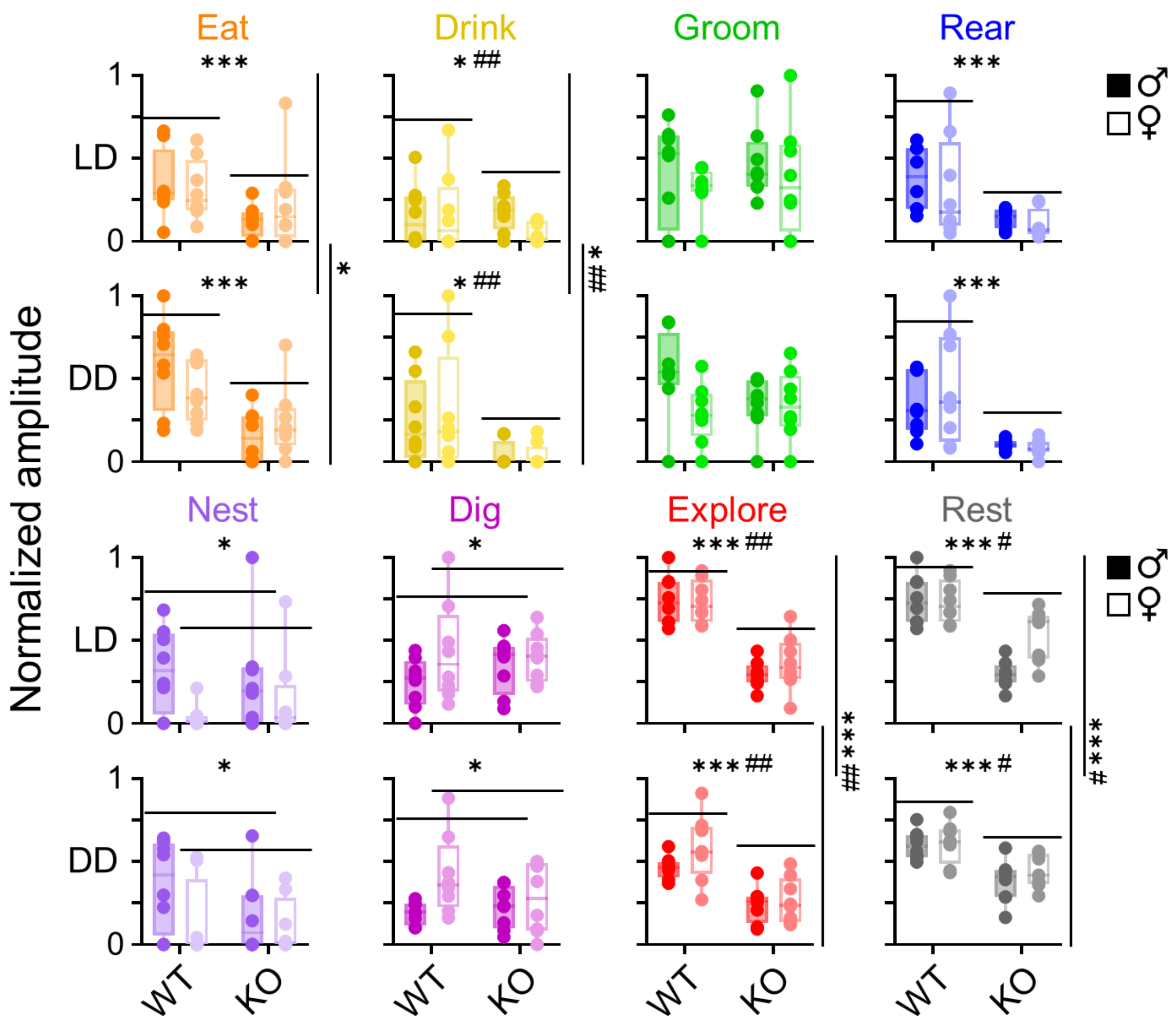


**Supplementary Figure 3. Amplitudes of behavioral rhythms differ by genotype, sex, and light cycle.** Rhythm amplitudes (in arbitrary units) for each behavior in individual wild-type and *Vip*$^{-/-}$ male (dark circles) and female (light circles) mice recorded in a 12 h:12 h light:dark cycle (LD) and in constant darkness (DD). Boxes indicate 25th to 75th percentiles. For visualization, rhythm amplitudes were normalized within each behavior. Three-Way Repeated Measures ANOVA, p < 0.05; ** p < 0.01; *** p < 0.001. #, significant interaction between behavior rhythm amplitudes across genotype, sex, or light cycle, p < 0.05; ## p < 0.01.

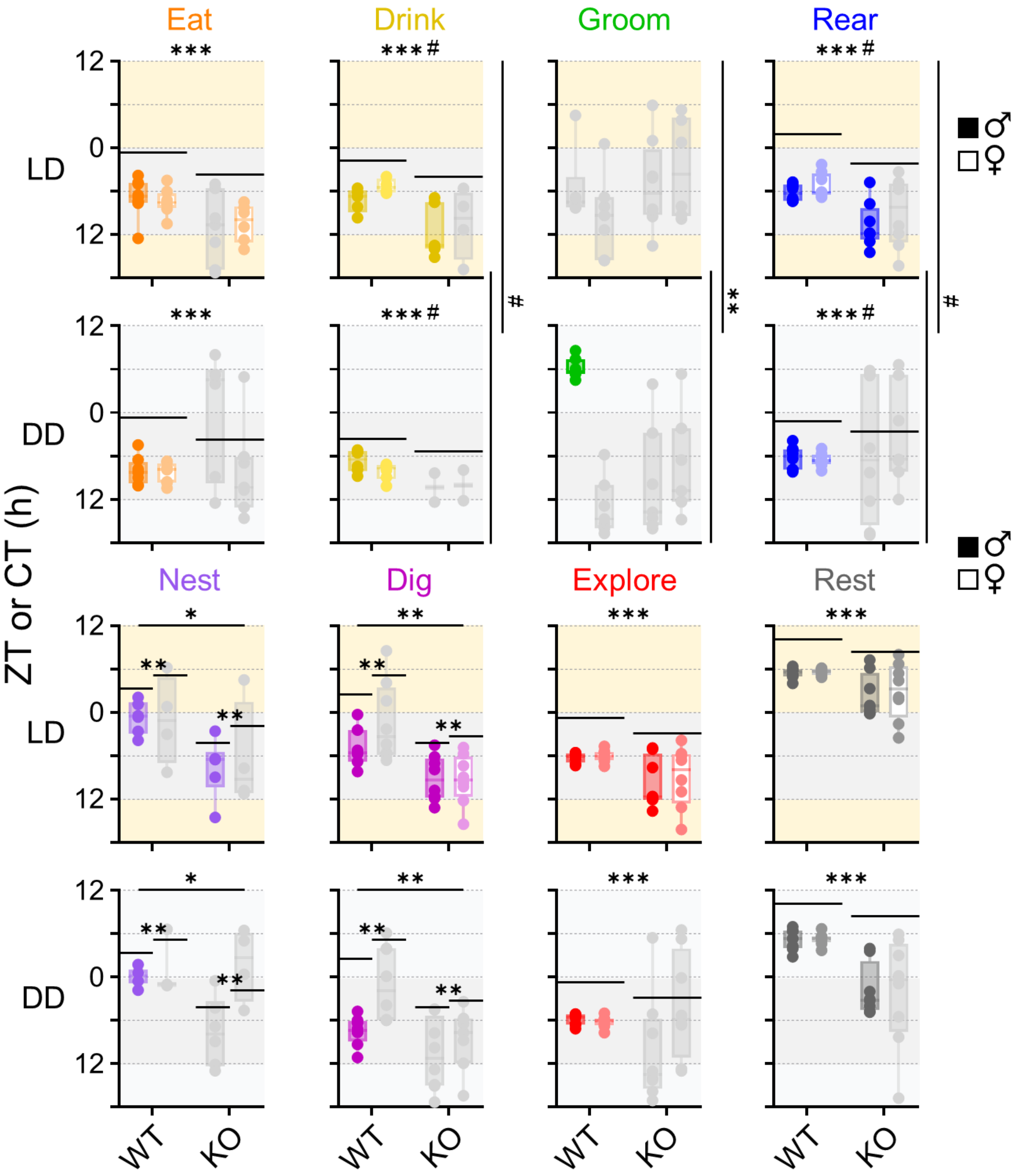


**Supplementary Figure 4. Peak times of behavioral rhythms differ by genotype, sex, and light cycle.** Peak times (in hours) for each behavior in individual wild-type and *Vip*$^{-/-}$ male (dark circles) and female (light circles) mice recorded in a 12 h:12 h light:dark cycle (LD; yellow and dark gray background) and in constant darkness (DD; light and dark gray background). Boxes indicate $25^{th}$ to $75^{th}$ percentiles. Gray dots and boxes depict rhythms that did not have significant clustering of peak times across mice (Rayleigh test, $p > 0.05$). Multi-Way Circular ANOVA, $p < 0.05$; ** $p < 0.01$; *** $p < 0.001$. #, significant interaction between peak times of behavioral rhythms across genotype, sex, or light cycle, $p < 0.05$. Peak times depicted as linear for visualization.

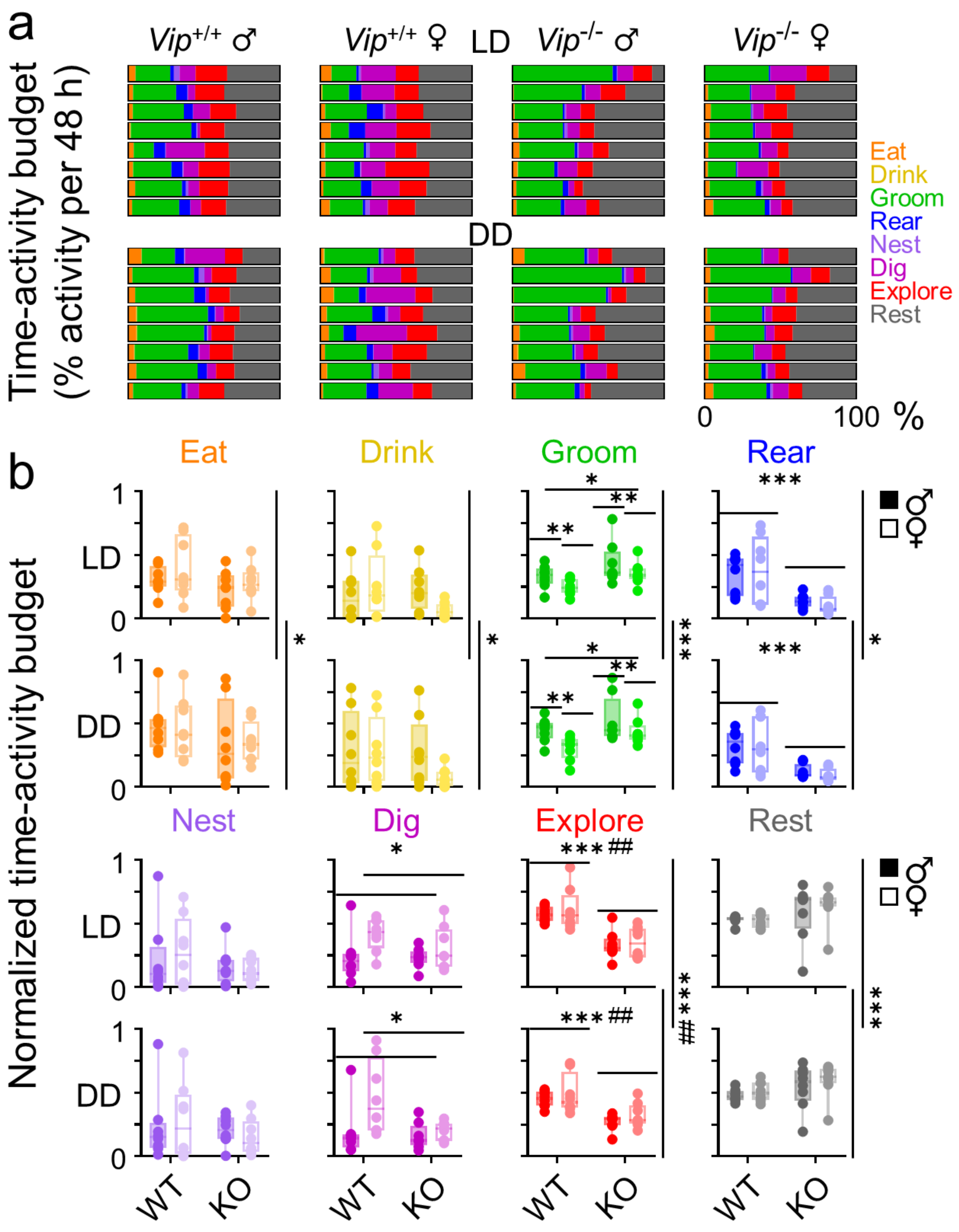


**Supplementary Figure 5. Behavior time budgets differ by genotype, sex, and light cycle. a)**

Time budgets (percentage of time spent in behavior across 48 h) of all behaviors for individual wild-type and *Vip*$^{-/-}$ male and female mice recorded in a 12 h:12 h light:dark cycle (LD) and in constant darkness (DD). **b)** Time budgets for each behavior in individual wild-type and *Vip*$^{-/-}$ male (dark circles) and female (light circles) mice in LD and in DD. Boxes indicate 25$^{th}$ to 75$^{th}$ percentiles. For visualization, time budgets were normalized within each behavior. Three-Way Repeated Measures ANOVA, $p < 0.05$; ** $p < 0.01$; *** $p < 0.001$. #, significant interaction between time budgets across genotype, sex, or light cycle, $p < 0.05$; ##, $p < 0.05$.

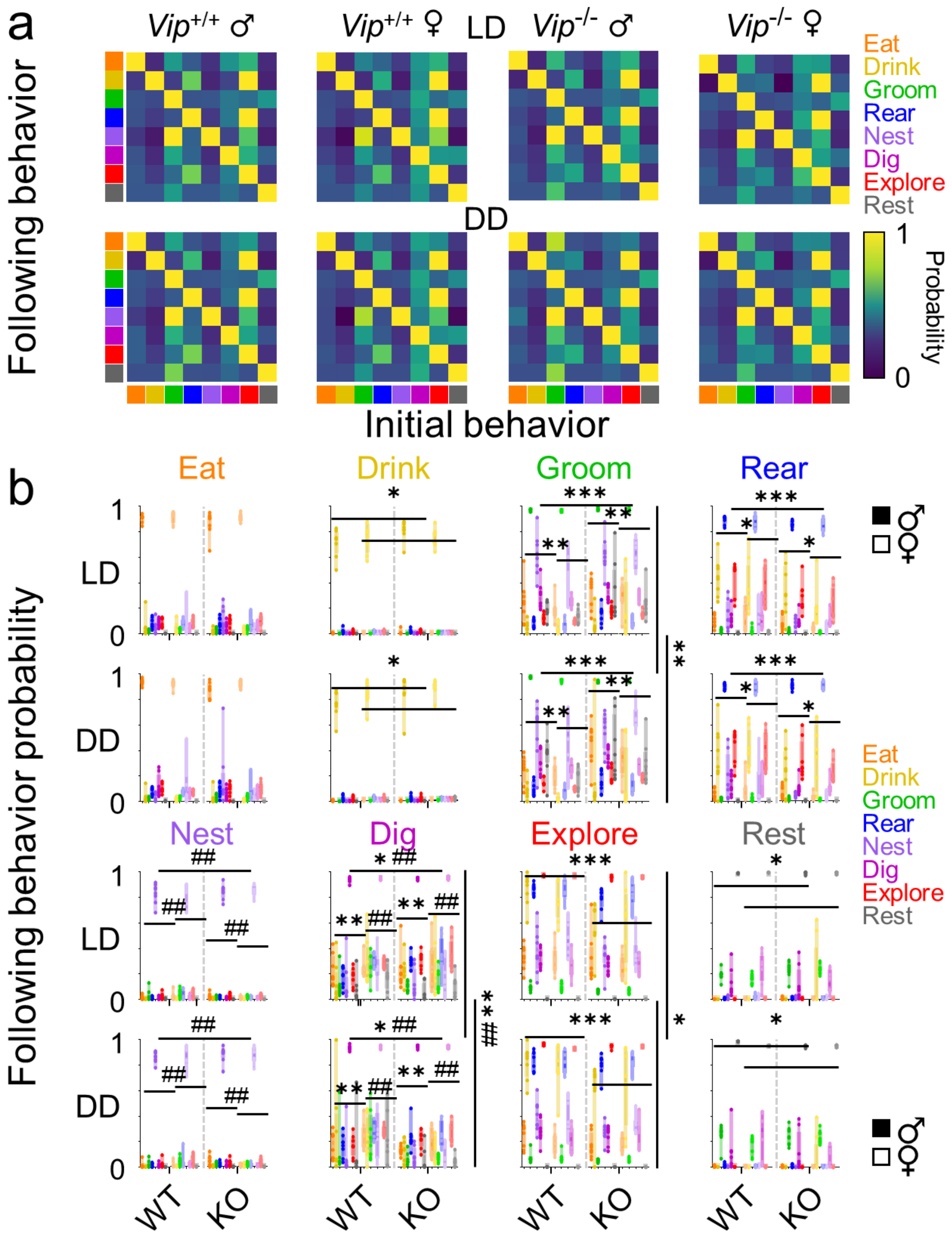


**Supplementary Figure 6. Behavioral state transitions differ by genotype, sex, and light cycle. a)** Transition matrices depicting transition scores (defined as the probability, from 0 to 1, that a given "following behavior" will follow an "initial behavior" within 10 seconds) averaged across mice of a given genotype and sex (n = 8 mice per condition) recorded in a 12 h:12 h light:dark (LD) cycle and in constant darkness (DD). Warmer colors indicate a higher probability of transitioning into that behavior. **b)** Transition probabilities for each initial behavior in individual wild-type and *Vip*$^{-/-}$ male (dark circles) and female (light circles) mice in LD and in DD. Boxes indicate 25$^{th}$ to 75$^{th}$ percentiles. Colored circles and bars within each "initial behavior" plot depict the transition probability for each "following behavior." Three-Way ANOVA, $p < 0.05$; **, $p < 0.01$, ***, $p < 0.001$. ##, significant interaction between transition probabilities across genotype, sex, or light cycle, $p < 0.01$.
